# Establishing a preclinical chronic wound model to characterise the pathophysiology of snakebite envenoming

**DOI:** 10.1101/2025.01.08.631785

**Authors:** Charlotte A Dawson, Pruistinne Harijanto, Yonlada Nawilaijaroen, Nicholas R. Casewell, Jenna L Cash

## Abstract

Snakebite claims 138,000 lives a year with an additional 400,000 patients left permanently disabled or disfigured1. Morbidity following envenoming includes the development of chronic wounds around the bite site. The understanding of the underlying pathophysiology of chronic snakebite wounds has been severely limited by the historical reliance on a preclinical model that only captures acute local envenoming pathology. Through the application of three medically important snake venoms (Echis ocellatus, Bothrops atrox and Naja nigricollis) to a recently developed preclinical model of chronic wounds, we have been able to characterise key features of venom wounds. We have been able to show that venom wounds share consistencies with non-venom induced preclinical wounds, and also display unique characteristics such as extracellular matrix degradation and eosinophilic infiltrate. This model will not only serve to increase our understanding the underlying pathophysiology of venom wounds, but will also provide a platform for exploring therapeutic interventions to reduce or resolve snakebite wounds.

## Introduction

Across the remote rural tropics, snakebite claims 138,000 lives a year^1^. In addition to this mortality, upwards of 400,000 snakebite patients annually are thought to be left permanently disabled or disfigured^1^. Morbidity following envenoming includes the development of chronic wounds around the bite site, and whilst data on the prevalence of these wounds is scarce, there are case studies documenting wounds persisting for years^2^.

Venom composition varies according to snake species, but is thought to typically induce hypoxia, inflammation, reactive oxygen species (ROS) and damage-associated molecular pattern (DAMP) release in the tissues surrounding the bite site^3,4^. The relative contribution and generality of these processes to the pathophysiology of local pathology remains poorly defined. Research into the chronic sequelae of local envenoming has been severely hampered by the field’s reliance on a local envenoming model that lacks a chronic dimension^5^. The established ‘minimum necrotic dose’ model only extends for 72-hours and was developed primarily as a tool to evaluate therapeutics rather than understanding venom pathology^5^.

Here, we establish a novel preclinical chronic wound model by delivering medically important venoms into the murine skin at doses previously demonstrated to induce dermal necrosis in the 72-hour envenoming model ^6,7,8^. Methodology for generating venom-induced chronic wounds was adapted from a recently described, novel preclinical model of chronic wounds in which mercaptosuccinic acid is used to induced wounds through the elevation of ROS^9^. Three compositionally and taxonomically diverse snake venoms (*Echis ocellatus, Bothrops atrox* and *Naja nigricollis*) were assessed to account for interspecific venom toxin variation (fig 1A). Briefly, venoms (*E. ocellatus*, 39 μg, *B. atrox*, 30 μg, or *N. nigricollis* 32, μg) were administered into 2 mm cutaneous wounds made into the depilated dorsal skin of male C57Bl6/J mice (n=6 per venom) (fig 1B). Venoms were delivered within a 30% pluronic hydrogel that sets at body temperature to ensure local envenoming without systemic effects. All animals received subcutaneous analgesia (buprenorphine at 0.1 mg/kg). Following venom delivery, wounds were covered with a transparent Tegaderm dressing to promote wound formation. Skin immediately around the dressing was preemptively treated with Bepanthen ointment to prevent irritation. Animals were monitored daily for 7 days to assess the development of local pathology at a macroscopic level (fig 2C) and to promptly identify and intervene, should PDSL be identified.

**Figure 1.**
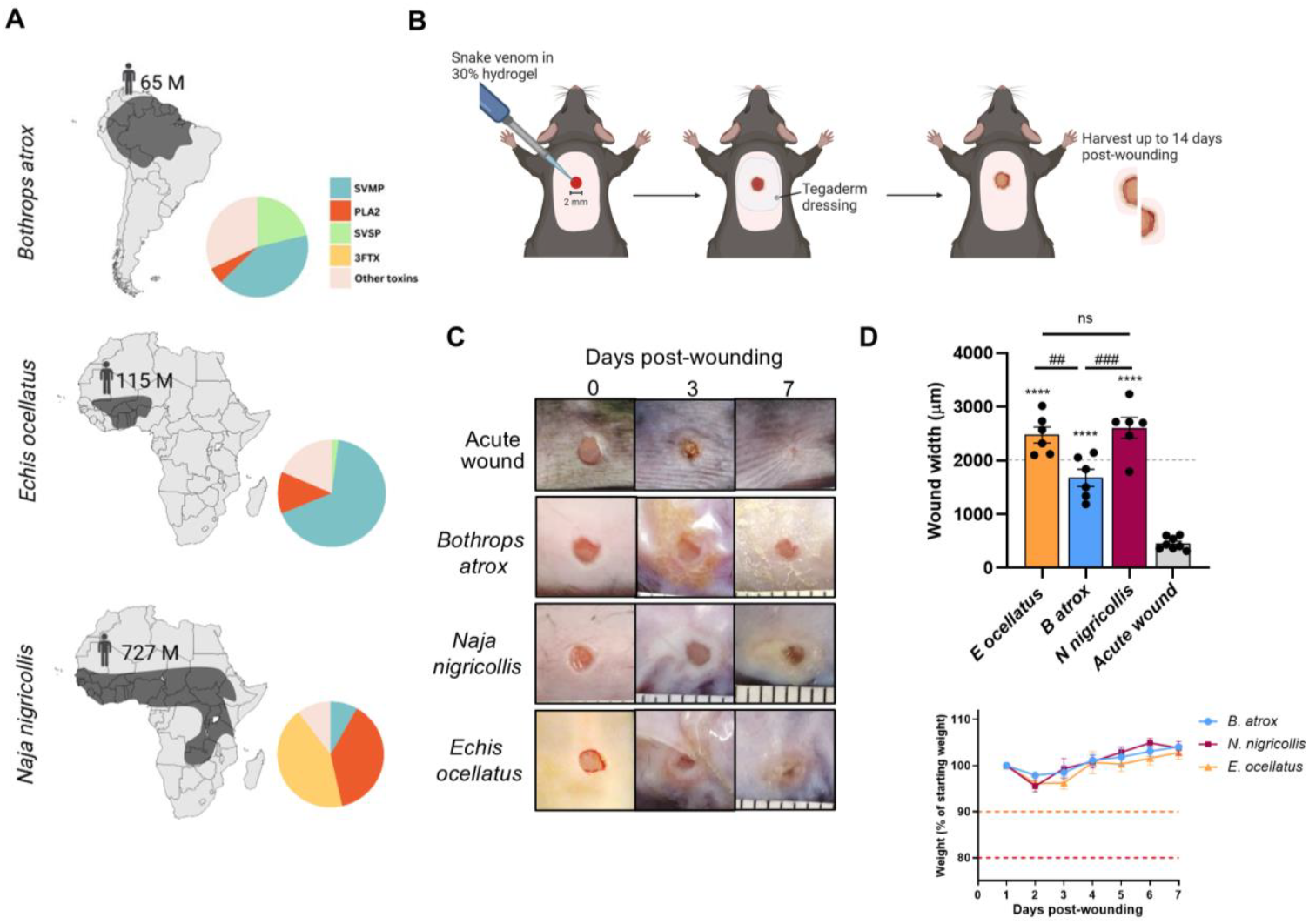
Development of venom-induced chronic wound model. **A** Geographic distributions of three medically important snake species from South America and sub-Saharan Africa (*B. atrox, E. ocellatus and N. nigricollis*) and the estimated human population in millions (M) at risk of envenomation, according to the WHO. Venom composition of these species, focussing on the more pathologically relevant toxins; Snake venom metalloproteinases (SVMP), phospholipase A2s (PLA2), Snake venom serine proteases (SVSP) and three-finger toxins (3FTX). **B** Schematic of experimental workflow. **B-C:** A single 2 mm excisional wound was made to depilated dorsal skin. *Echis ocellatus* (39 μg), *Bothrops atrox* (30 μ g) or *Naja nigricollis* (32 μ g) venom in 30% hydrogel carrier were topically administered to the wound immediately after injury. Wounds were covered with Tegaderm dressing and monitored over 7 days post-envenoming. **D Top:** Wound width 7 days post-envenoming in comparison to acute wound healing. ****, p<0.0001 relative to acute wounds, ##, p<0.01; ###, p<0.001 relative to 8 atrox chronic wounds by 1-way ANOVA. n=6 per venom and n=8 for acute wound. Grey dotted line indicates wound width at day 0. **Bottom:** Body weight monitoring reveals minimal weight loss following wound envenomation. No systemic effects were noted and survival was 100%. Orange dotted line = increased monitoring threshold, red dotted line = cull threshold

**Figure 2.**
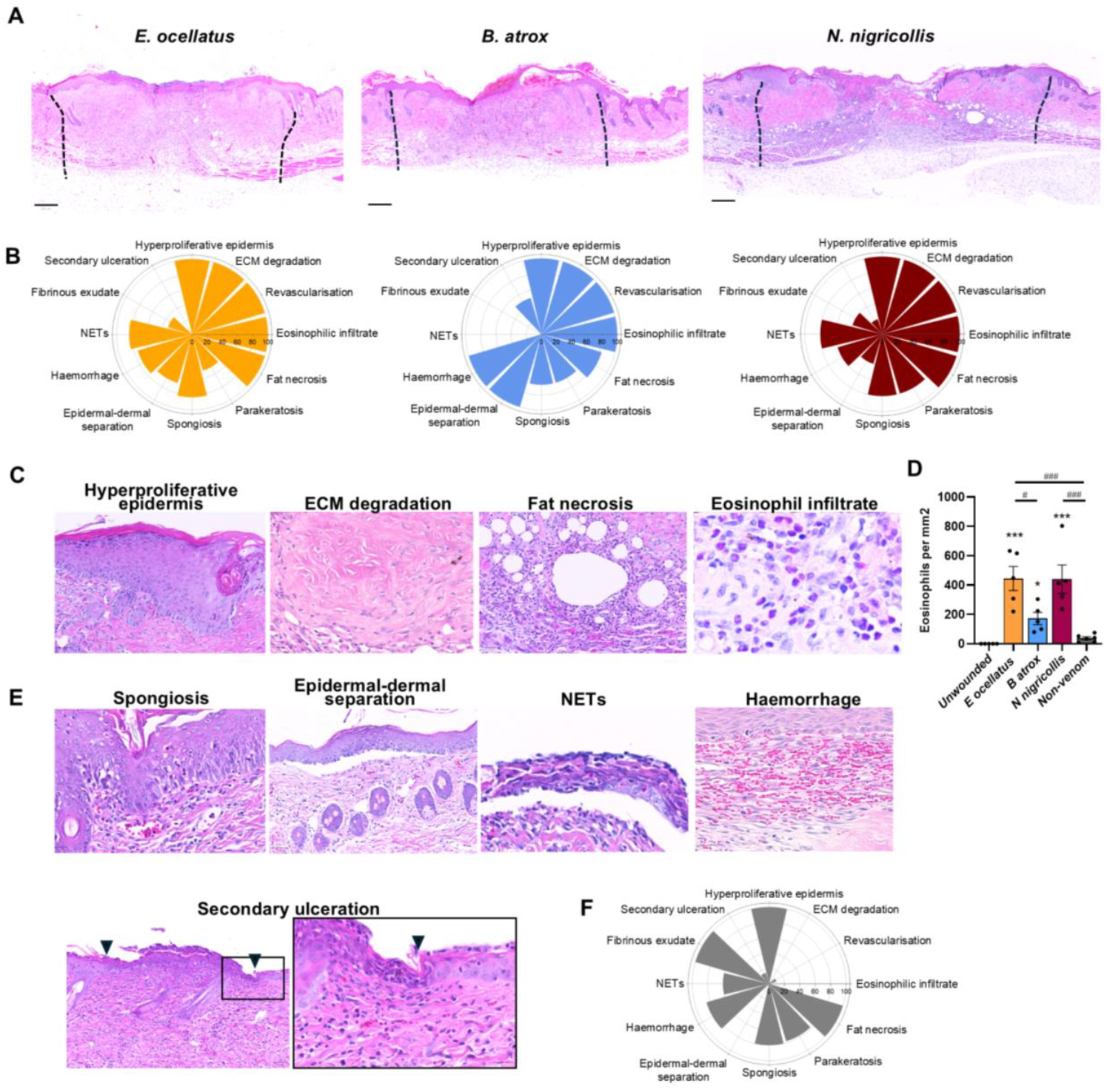
Histomorphological characterisation of venom-induced wounds reveals distinct characteristics compared to non-venom wounds. **A** Representative H&E-stained wound mid-sections 7 days post-envenoming with 39 μ g *Echis ocellatus*, 32 μg *Naja nigricollis* and 22 μ g *Bothrops atrox venom*. Scale bars indicate 250 pm. Black dotted lines indicate wound margins and represent the border between venom-induced extracellular matrix (ECM) degradation and typical ECM. **B** Rose plots indicating percentage incidence of histomorphological features identified in H&E-stained wound sections 7 days post-envenoming. **C** Core histomorphological features observed in all wounds regardless of venom type; hyperproliferative epidermis, ECM degradation, fat necrosis and eosinophilic infiltrate. **D** Quantification of eosinophilic infiltrate. 1 way ANOVA, *, p<0.05; ***, p<0.001 relative to unwounded skin. #, p<0.01; ###, p<0.001 relative to non-venom chronic wounds 7 days post-injury. **E** Representative images of other commonly observed histomorphological features. Black triangles mark borders of secondary ulceration. **F** Rose plot indicating percentage incidence of histomorphological features in chronic wounds induced by locally raising oxidative stress in aged animals using mercaptosuccinic acid, n = 6 per group. NETs, neutrophil extracellular traps

Venom specific differences were observed immediately after venom delivery, with the viperid venoms (*B. atrox* and *E. ocellatus*) causing rapid onset localised haemorrhage, whilst wounds produced by envenoming from the elapid *N. nigricollis* resulted in clear straw-coloured exudate without evidence of local haemorrhage. In all venoms a white area around the initial was observable within 6-hours of dosing. Wound widths measured 7 days post-injury revealed that all venoms induced wounds that were significantly larger compared to non-venom induced acute wounds (fig 1D).

Animal welfare was monitored throughout the study duration (fig 1E). Animals were able to maintain, and increase, their body weights indicating that wounds were well tolerated. Irritation of the skin surrounding the dressing area peaked at days 3 – 4 post wounding, ranging from 60 – 80% of animals per group depending on venom. Whilst the proportion of animals displaying irritation was greater than the previously described non-venom chronic wound model^9^, topical application of bepanthen was sufficient to manage irritation.

On day 7 post wound, animals were humanely euthanised via rising CO_2_, and wounds harvested. Wound mid-sections were obtained for haematoxylin and eosin staining to enable visualisation and quantification of histomorphological characteristics (fig 2A). All wounds showed several common features, including hyperproliferative epidermis, degradation of the extracellular matrix (ECM), and evidence of revascularisation (fig 2B-C). Fat necrosis was also seen across all *E. ocellatus* and *N. nigricollis* wounds, and in 83% of wounds induced with *B. atrox* (fig 2B-C). All wounds showed a significant infiltration of eosinophils (fig 2B-D). Quantification of the number of cell per mm^2^ found that a greater eosinophil infiltrate was seen in *E. ocellatus* and *N. nigricollis* venom compared to *B. atrox* (444.8 cell/mm^2^ and 439.8 cells/mm^2^ respectively vs. 172 cells/mm^2^), however, all venom wounds had significantly more eosinophils that in non-venom induced wounds (37.3 cells/mm^2^)

Venom-specific variation in additional wound characteristics was also observed (fig 2E). The viperid venoms, *B. atrox* and *E. ocellatus*, had haemorrhage present in more wounds (100% and 75% respectively) compared to *N. nigricollis* (60%) at 7 days post-envenoming. Epidermal-dermal separation was more prevalent in response to vipers, *E. ocellatus* and *B. atrox* (67% and 100% respectively, vs 40% in *N. nigricollis* wounds), conversely, wounds caused by *N. nigricollis* had greater incidence of parakeratosis (80%) compared to either *B. atrox* or *E. ocellatus* (67% and 50%). Secondary ulceration was observed in wounds caused by *B. atrox* (50%) and N. *nigricollis* (20%) but not with *E. ocellatus*.

Lastly, histomorphological features of venom wounds were compared against those seen in non-venom chronic wounds (fig 2F). Whilst both models showed hyperproliferative epidermis, fat necrosis, spongiosis and haemorrhage, neutrophil extracellular traps (NETs), there were several distinct features. The observation of ECM degradation, eosinophilic infiltrate, epidermal-dermal separation and revascularisation was only recorded in venom wounds, suggesting these features may be unique to envenoming (Fig.2B,F).

Understanding the underlying pathophysiology of venom-induced chronic wounds and determining the commonalities and differences of these compared to wounds of other causations, is essential for developing strategies to alleviate the burden of snakebite morbidity. Cytotoxic envenomings contribute significantly to the annual morbidity of snakebite, of which venom-induced chronic wounds are a poorly understood component. To better understand the underlying wound aetiology, we applied three medically important venoms to a novel chronic wound model previously described by Nawilaijaroen *et al* (9). The application of venom to this model generated wounds that showed a failure to resolve across 7 days, compared to acute, non-venom induced wounds. Venom-induced wounds showed several features consistent with non-venom wounds, such as fat necrosis, parakeratosis, haemorrhage and NETs. However, there were several features seen only in the venom model, including ECM degradation, revascularisation and a significant eosinophilic infiltrate.

The insight gained into the underlying mechanisms of venom induced wounds also opens new therapeutic opportunities. Current snakebite therapeutics focus solely on inhibiting the action of venom toxins, however intervening in downstream host processes that drive wound formation and exacerbate the severity of local envenoming, or that stimulate the healing of wounds is an unexploited, yet potentially efficacious, therapeutic strategy to mitigate the devastating, often lifelong, consequences of snakebite. Future use of this novel venom-induced chronic wound model will enable robust evaluation of such therapeutics with the goal of mitigating the severe, often life-changing, effects of tropical snakebite.

## Ethics statement

All experiments were conducted with approval from the University of Edinburgh Local Ethical Review Committee and in accordance with the UK Home Office regulations (Guidance on the Operation of Animals, Scientific Procedures Act, 1986) under PPL PP2067887.

## Acknowledgements

We thank the Liverpool School of Tropical Medicine for the funding that enabled this work. We also thank Paul Rowley and Edouard Crittenden for the maintenance and husbandry of the snake collection and the provision of venom samples held at LSTM. At the University of Edinburgh, we thank the Biological veterinary services for their support with animal experiments, and SURF for support with histological services.

## Author contributions

CD: Conceptualisation, Methodology, Investigation, Writing – Original Draft, Project administration, Funding acquisition PH: Investigation. YN: Investigation. NC: Writing – Review & Editing, Resources. JC: Conceptualisation, Methodology, Formal analysis, Investigation, Resources, Writing – Original draft, Visualization, Supervision, Project administration.

## References

1. Gutiérrez JM, Calvete JJ, Habib AG, Harrison RA, Williams DJ, Warrell DA. Snakebite envenoming. Nat Rev Dis Primer. 2017 Sep 14;3(1):1–21.

2. Jayawardana S, Gnanathasan A, Arambepola C, Chang T. Chronic Musculoskeletal Disabilities following Snake Envenoming in Sri Lanka: A Population-Based Study. De Silva J, editor. PLoS Negl Trop Dis. 2016 Nov 4;10(11):e0005103.

3. Rucavado, A., Nicolau, C. A., Escalante, T., Kim, J., Herrera, C., Gutierrez, J. M., Fox, J. W. Viperid Envenomation Wound Exudate Contributes to Increased Vascular Permeability via a DAMPs/TLR-4 Mediated Pathway. Toxins. 2016. 8 (12).

4. Katkar, G. D., Sundaram, M. S., NaveennKumar, S. K., Swethakumar, B., Sharma, R. D., Paul, M., Vishalakshi, G. J., Devaraja, D., Girish, K. S., Kemparaju, K. NETosis and lack of DNAase activity are key factors in Echis carinatus venom-induced tissue destruction. Nat Comms. 2016. 7.

5. Theakston RDG, Reid HA. Development of simple standard assay procedures for the characterization of snake venoms. Bull. World Health Organ. 1983. 61 (6). 949–56.

6. Hall, SR., Rasmussen, SA., Crittenden, E., Dawson, CA., Bartlett, KE., Westhorpe, AP., Albulescu L-O., Kool, J., Gutierrez, JM., Casewell, NR. Repurposed drugs and their combinations prevents morbidity-inducing dermonecrosis caused by diverse cytotoxic snake venoms. Nature Communications. 2023.14. 7812. doi: 10.1038/s41467-023-43510-w

7. Ahmadi, S., Pachis, ST., Kalohgeropoulos, K., McGeoghan, F., Canbay, V., Hall, SR., Crittenden, EP., Dawson., CA., Bartlett, KE., Gutierrez, JM., Casewell, NR., auf dem Keller, U., Lausten, AH. Proteomics and histological assessment of an organotypic model of human skin following exposure to Naja nigricollis venom. Toxicon. 2022. 220. 106955.

8. Santos Barreto GNL, De Oliveira SS, Dos Anjos IV, Chalkidis HDM, Mourão RHV, Moura-da-Silva AM, et al. Experimental Bothrops atrox envenomation: Efficacy of antivenom therapy and the combination of Bothrops antivenom with dexamethasone. Calvete J, editor. PLoS Negl Trop Dis. 2017. 11(3). e0005458.

9. Nawilaijaroen, Y., Rocliffe, H., Austin-Williams, S., Krilis, G., Pellicoro, A., Zhou, K., Ji, Y., Bain, C. a., Kilpartrick, A. M., Chen, Y., Biswas, A., Crichton, A., Huang, Z., Forbes, S. J., Caporali, A., Cash, J. L. MC1R reduces scarring and rescures stalled healing in a preclinical chronic wound model. BioRxiv. 2022. 11.30.518516

